# Lightweight open-source fine-tuning of SAM2 enables domain-specific microscopy segmentation

**DOI:** 10.1101/2025.11.08.687405

**Authors:** Etash Bhat, Sharvaa Selvan, Sonia Okekenwa, Zareh Dechkounian, Vivian Lin, Mayu Nakano, Mitul Saha, Yuyu Song

## Abstract

Quantitation of structures is a critical step in analyzing images. Automated segmentation of biological samples remains a central challenge in microscopy, where variations in signal/noise, intensity, texture, and edges hinder accurate delineation of cellular and tissue structures. Adaptations of foundation models such as Segment Anything Model (SAM) remain computationally intensive and require large training datasets. Here, we introduce a lightweight, open-source Google Colab pipeline that enables efficient fine-tuning of SAM2 on domain-specific datasets without additional architectural layers or specialized hardware. By coupling mask-decoder fine-tuning with biologically informed post-processing, our framework achieves robust segmentation across diverse imaging modalities. Applied to hippocampal segmentation in brain images and single-cell segmentation in cell images, fine-tuned SAM2 demonstrates substantial gains of accuracy relative to basic SAM2 and matches leading tools. This work establishes a scalable and accessible paradigm for domain-specific adaptations of SAM2 in microscopy, lowering computational and data barriers to advanced image segmentation.

**Highlights:**

- **Lightweight fine-tuning** with no added architectural complexity.
- **High segmentation accuracy** (Dice/Jaccard scores) achieved with small datasets.
- **Cross-domain generalization** across tissue and cell imaging with biologically informed post-processing.
- **Comparable or superior performance** to widely used tools (Cellpose, Imaris, ilastik) at substantially lower computational cost.
- **Open-source and executable in a single Colab notebook**, ensuring reproducibility and accessibility for non-computational users.
- **Turnkey adaptability**, allowing researchers to transform raw microscopy data into fine-tuned SAM2 models with minimal input.

## Main

Microscopic visualization of cellular and tissue structures is critical for quantifying biomarkers and understanding biological function^1^. A wide range of microscopy techniques, from confocal microscopy for high-resolution optical sections^2^ of cell samples to virtual slide microscopes^3-5^ well-suited for larger-scale tissue morphology, has made automated segmentation of regions of interest (ROIs) an essential yet challenging task in image analysis^6^. Several open source tools, including ImageJ^7,8^, Imaris^9^, and ITK-SNAP^10,11^, provide segmentation engines that use classical image analysis methods to extract masks that correspond to ROIs. However, these traditional methods often demonstrate low performance in images that have low contrast, high noise, complex texture, or out-of-focus cells by generating masks that retain artifacts, fragment single cells, and fail to capture cell boundaries^12,13^.

Machine learning methods such as Cellpose^14,15^, ilastik^16^, and CellProfiler^17^ improve ROI segmentation but remain either overly specialized or labor-intensive to prepare for each imaging modality. More recently, foundation models such as the Segment Anything Model^18^ (SAM), and its enhanced model SAM2^19^, have shown impressive generalization. However, the default SAM relies on uniform grid prompting^20^, which leads to fragmented masks in images with non-uniform cell densities and poor delineation of tissue structures where ROIs are restricted to only a subset of the field.

Adaptations such as μSAM^21^ (new decoder + iterative prompting) and CellSAM^20^ (transformer-based prompting) address these challenges; however, their reliance on additional transformers or convolutional layers makes them computationally demanding and less scalable as fine-tuning strategies for laboratories seeking to adapt SAM to their own datasets. Therefore, there is still an unmet need for a computationally elegant solution that 1) enables SAM2 fine-tuning without additional architectural complexity, 2) can be trained on small, domain-specific datasets, and 3) remains accessible to researchers without specialized hardware. Here, we introduce a lightweight Google Colab pipeline for fine-tuning SAM2 using modest, curated microscopy datasets (Figure 1a). By combining targeted mask decoder fine-tuning with algorithmic mask optimization, our framework provides a scalable and open-source approach for adapting SAM2 to diverse applications.

**Figure 1.**
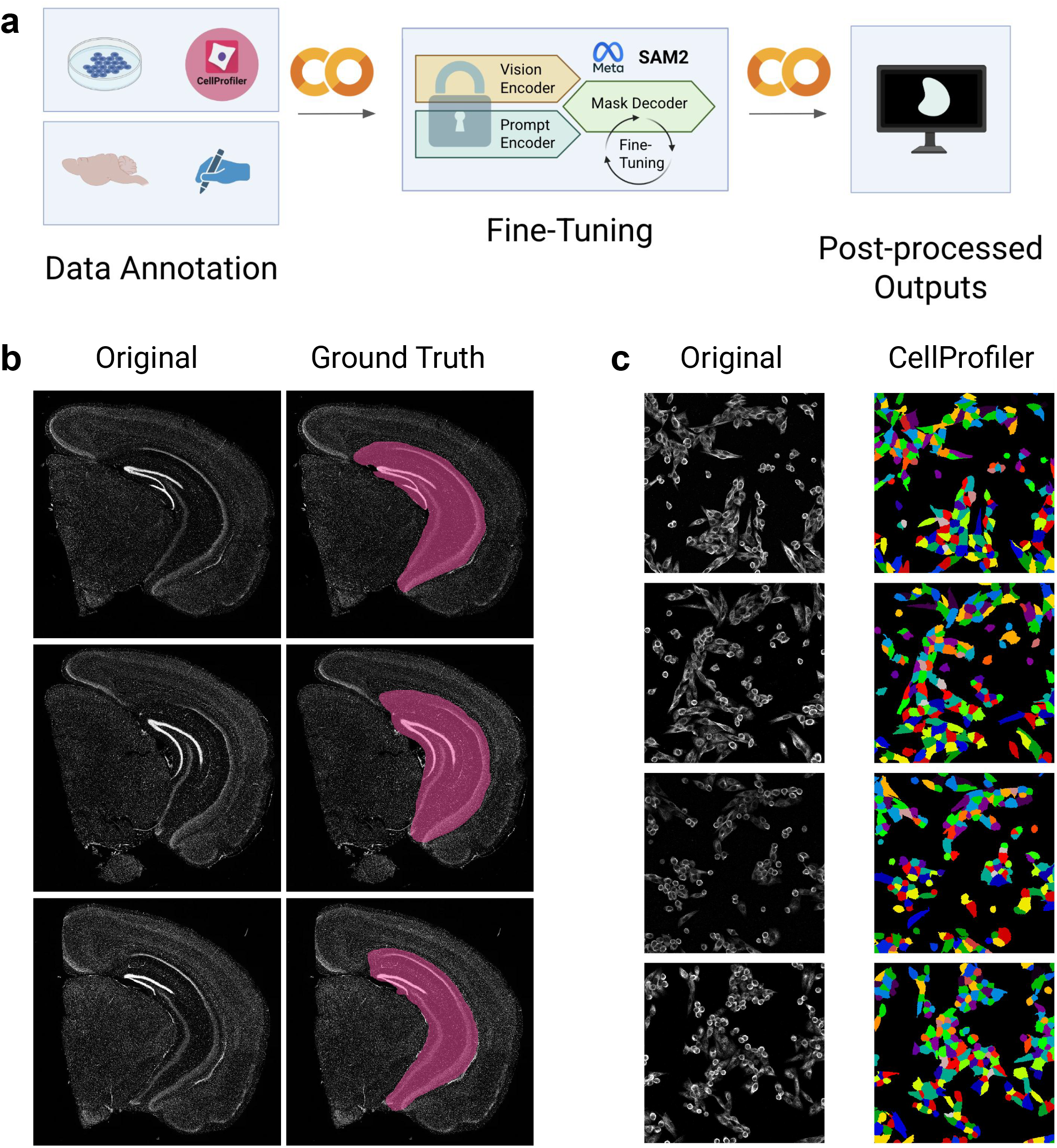
Lightweight Google Colab pipeline enables domain-specific SAM2 adaptations for image segmentations. Fine-tuning the SAM2 mask decoder with a compact dataset, paired with output post-processing, establishes a scalable and accessible strategy for adapting the SAM2 to diverse microscopy applications. a) Images collected by individual laboratories can be annotated by software (cell images) or manually (more specialized applications) to train the SAM2 mask decoder while keeping other computational components frozen. Biologically constrained algorithms can refine the output of the fine-tuned SAM2, achieving high segmentation quality across brain and cell imaging domains. b) Mouse brain images with manually annotated hippocampi and c) cell images annotated by CellProfiler serve as training data for SAM2 fine-tuning.

We demonstrate the functionality of our pipeline by the fine-tuning of SAM2 with 54 Hoechst-stained mouse brain images (and corresponding ground-truth masks) to accurately segment hippocampi. This represents the first optimization of a large-scale foundational model for mouse brain image segmentation (Figure 1b). We further develop a pixel-based post-processing algorithm that can refine the outputs of the fine-tuned SAM2 by reliably selecting the most accurate mask among the top-confidence predictions. To apply our pipeline to more rigorous segmentation tasks, we fine-tune SAM2 with 59 tubulin-stained Chinese Hamster Ovary (CHO) cell images and corresponding masks annotated automatically using CellProfiler (Figure 1c). For algorithmic mask optimization, we use a novel post-processing algorithm involving corresponding Hoechst stains of the images. Our results demonstrate excellent performance compared with current leading standards, like Imaris, Ilastik, Cellpose, and Otsu thresholding, at a low computation cost.

## Results

To validate our lightweight pipeline of fine-tuning and post-optimization, we evaluated performance across increasingly complex segmentation tasks. We first test the approach on hippocampus segmentation from mouse brain images with nuclear Hoechst staining to extract a single, continuous ROI. We then evaluate our pipeline on the more rigorous task of single-cell segmentation, using parameter optimization and Hoechst-guided post-processing to enhance the fine-tuning output. Finally, we compare our lightweight pipeline with various widely used segmentation software and other SAM adaptations.

### Proof-of-concept: hippocampus segmentation

We first evaluated our lightweight pipeline on the task of segmenting hippocampi from Hoechst-stained mouse brain images. To that end, we fine-tuned the SAM2 mask decoder^19^ with 54 images paired with ground truth and evaluated the highest-confidence output for each image on a held-out test set of 24 images. Illustrating our approach, Figure 2a shows representative images of ground truth, default outputs, fine-tuned outputs, and the highest-confidence outputs in the four leftmost columns with final refined output in the right-hand column. Performance of the fine-tuned SAM2 was compared with the default SAM2 using two measures of accuracy^22^ relative to the ground truth: Dice score^23^ (overlap similarity) and Jaccard^24^ index (intersection over union). We expected improvement compared to the default SAM2 on both measures, even without the addition of deep learning architectures. Indeed, fine-tuning improved performance from 0.091 to 0.646 (mean Jaccard) and from 0.157 to 0.686 (mean Dice), as shown in Figure 2b.

**Figure 2.**
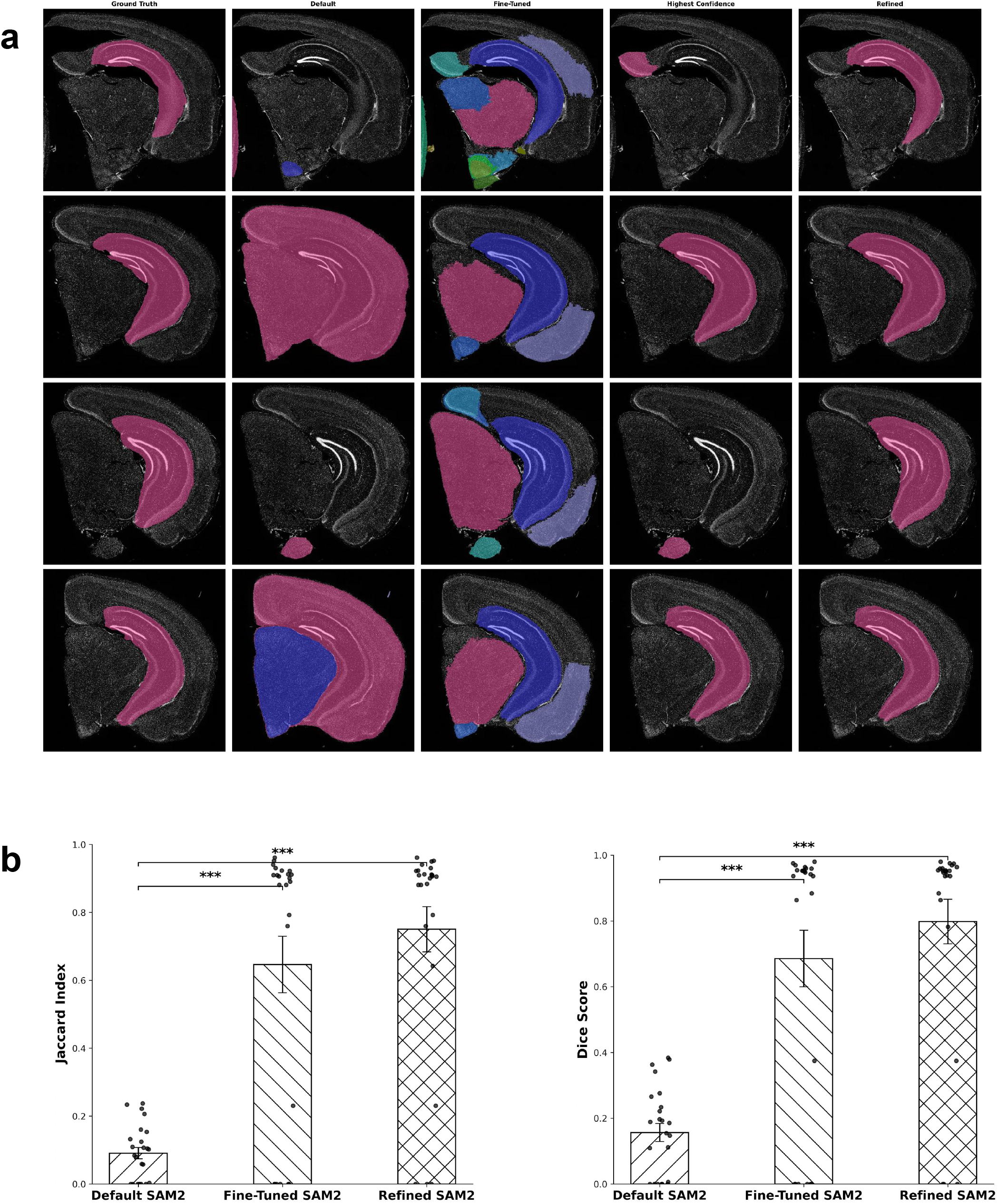
Accurate hippocampus segmentation from mouse brain images via fine-tuned SAM2 with pixel-based post-processing. Our pipeline achieves accurate hippocampus segmentation by fine-tuning the SAM2 mask decoder with annotated mouse brain images and refining outputs using a pixel-intensity criterion, informed by the high cell density of the hippocampus. a) Representative mouse brain images and intermediate pipeline outputs, illustrating the progression from Base SAM2 outputs to fine-tuned outputs, highest-confidence masks, and biologically refined outputs, compared against ground truth annotations. b) Dice scores and Jaccard indices were measured against the ground truth for the Base SAM2 outputs, fine-tuned outputs, and biologically refined outputs. Error bars represent standard error of the mean across independent image samples.

Though fine-tuning improves the segmentation of the mouse brain images, the overall performance is dispersed across images (Figure 2b). Notably, highest-confidence masks outputted by the fine-tuned SAM2 fall into two categories: either they are precisely correct, or they miss the region entirely. This is because the different anatomical regions of the mouse brain in these images share, to a certain degree, similar texture, color, and shape — all of which are cues that the SAM2 mask decoder uses to carve out the correct mask. Therefore, incorrect masks segment tissue surrounding the hippocampus, although they use semantic cues that mimic those of the hippocampus. However, we noticed that the correct mask is always generated by our model, even if it is not the highest-confidence mask. Because our method of fine-tuning SAM2 on a modest dataset may lead to overfitting, we hypothesized that the consistent presence of the correct mask can be leveraged through post-processing.

### Post-processing improves hippocampus segmentation

Building on this insight, our lightweight pipeline integrates algorithmic post-processing of the fine-tuned SAM2 output based on relevant biological features to generate the final ROI segmentation, hereafter called the “refined SAM2”. In the case of mouse hippocampus segmentation, we used dentate gyrus as a hallmark, which has the densest nuclei stained by Hoechst showing the highest pixel intensities. Based on this observation, we developed a simple algorithm of identifying the 3 most confident mask predictions from the fine-tuned model and selecting the one with the highest pixel intensity. Representative images of the refined SAM2 outputs are shown on the far-right column of Figure 2a.

To evaluate the improvement in hippocampus segmentation of the refined SAM2, which combines mask-decoder fine-tuning with algorithmic post-processing, we computed the Dice score and Jaccard index in the same manner as before. Because the refined SAM2 is designed to improve the fine-tuned output with a biologically informed algorithm, we expected the accuracy measures to be substantially higher than those of the fine-tuned SAM2. The results confirmed this, with performance rising from 0.646 to 0.750 (mean Jaccard) and from 0.686 to 0.798 (mean Dice), as shown in the far-right bars of Figure 2b. Remarkably, most test images clustered around performance values near 0.9 for both metrics.

The drastic improvement underscores how modest fine-tuning, paired with algorithmic mask optimization, can achieve high segmentation quality and accuracy without the expansive datasets and heavy computational architecture of more resource-intensive approaches. Importantly, this case study demonstrates that simple segmentation tasks can benefit substantially from a lightweight pipeline. This case study also establishes a precedent for optimizing large-scale foundation models for the task of mouse brain image segmentation. Having established the effectiveness of our lightweight pipeline in brain tissue, we next turned to the more challenging problem of segmenting individual cells, where object heterogeneity, overlap, and noise make accurate instance segmentation considerably more demanding^25^.

### Single-cell segmentation with lightweight SAM2 fine-tuning

To evaluate our lightweight pipeline on the task of segmenting cells from tubulin-stained CHO cell images, we fine-tuned the SAM2 mask decoder with 59 images paired with automatically generated ground truth from CellProfiler. Naturally, we expected the performance, compared to manually segmented ground truth images, to improve relative to the Base SAM2 because of the domain-specific training. Surprisingly, on a held-out test set of 25 images, performance declined from 0.184 to 0.048 (mean Jaccard) and from 0.275 to 0.089 (mean Dice) for aggregate mask area: the fine-tuned SAM2 outputted only a small fraction of the correct masks (Figure 3). We compared the original, manually segmented, default SAM2 (base), and fine-tuned SAM2 segmented images. The results showed visually poor segmentation performance of the aggregate mask area, so we decided to first optimize the pipeline to constrain the total number of candidate masks before paring the masks down to the ideal set of single-cell segmentations.

**Figure 3.**
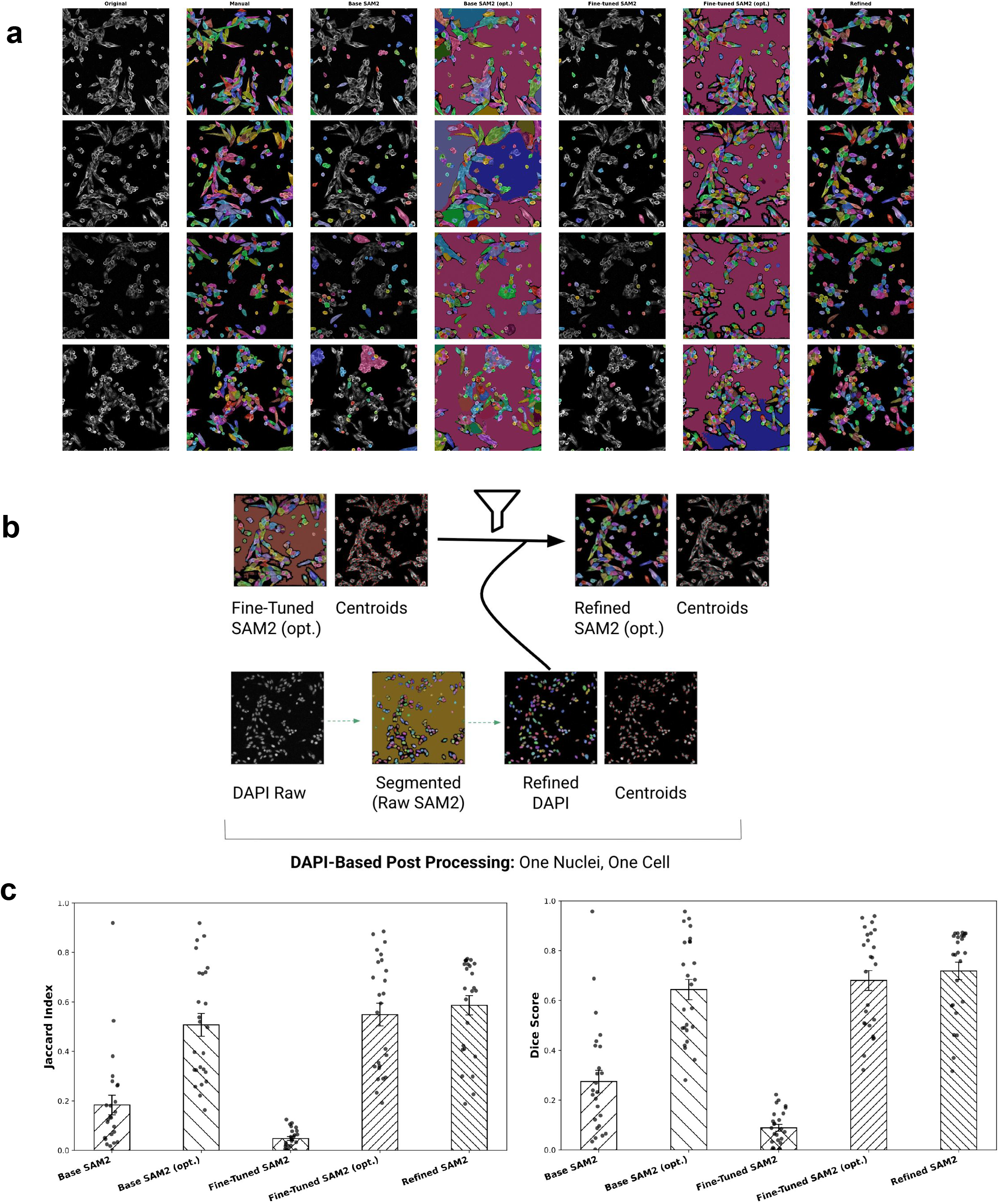
Nucleus-anchored refinement improves CHO cell segmentation from fine-tuned SAM2 outputs. To address CHO cell instance segmentation, we fine-tune the SAM2 with CellProfiler annotations and refine the output by enforcing a one-to-one correspondence with Hoechst nuclear masks. a) Representative CHO cell images and intermediate pipeline outputs, illustrating the progression from Base SAM2 outputs to fine-tuned outputs, parameter-optimized outputs, and biologically refined outputs, compared against manual ground truth annotations. b) Corresponding Hoechst masks for each image are segmented with the Base SAM2 and pruned of duplicates (Refined Hoechst) before the mask centers are used to filter the fine-tuned SAM2 output (Refined SAM2 opt.). See the Methods for the details. c) Dice scores and Jaccard indices were measured against the manually generated ground truth for each SAM2 iteration. Error bars represent standard error of the mean across independent image samples.

To understand why fine-tuning, unlike in the brain segmentation task, failed to immediately produce correct aggregate mask areas for any images, we investigated the SAM2 architecture.

The specific SAM2 mask outputs are governed by tunable parameters^19^ that control mask density and filtering (e.g. minimum mask region area, stability score). Aligning with our broader pipeline of the fine-tuning of the SAM2 with post-processing of the outputs, we carried out a simple parameter optimization pipeline on a validation subset. The best-performing configuration of parameters was selected for the fine-tuned SAM2, yielding an optimized version of the model (hereafter called the “optimized SAM2”). The optimized SAM2 was reevaluated, demonstrating improvement from 0.184 to 0.508 (mean Jaccard) and from 0.275 to 0.644 (mean Dice) for aggregate mask area compared to the Base SAM2 with default parameters (Figure 3c). Representative images of Base SAM2 and fine-tuned SAM2 with optimized parameters (Figure 3a, columns 4 and 6, respectively) and quantification (Figure 3c) demonstrate that SAM2 fine-tuning, paired with algorithmic parameter optimization, enables segmentation optimization without significant resources.

Despite the marked improvement of the optimized SAM2 compared to the Base SAM2 at segmenting aggregate area, the model output hundreds more masks than the ground truth: several cell regions contained two or three mask centers. This is because optimized SAM2 struggled to differentiate between cell boundaries, thus lumping adjacent cells into a single mask while also retaining the individual cells masks, leading to a tangle of overlapping masks. Importantly, however, we observed that the correct masks were always contained as the subset of the total output. Thus, the behavior of the optimized SAM2 model parallels that of the original fine-tuned model for hippocampus segmentation: the generated masks can be systematically parsed to isolate the correct subset. Building on our parameter optimization algorithm (see the Methods for details), we extended the algorithm based on biologically informed constraints to yield the final, accurate set of single-cell masks.

### Hoechst-based post-processing improves single-cell segmentation

Our lightweight post-optimization algorithm leverages the fact that each CHO cell has exactly one nucleus at a given plane of imaging. Using corresponding Hoechst nuclear stain, which is routinely used for cell culture imaging, for each image, the algorithm refines our optimized SAM2 by enforcing a one-to-one correspondence between nuclei (segmented by Base SAM2) and cell masks. Figure 3b highlights key steps in the algorithm with a representative image and its corresponding Hoechst stain. In evaluating this new model (“refined SAM2”), we expected a substantial increase in the accuracy of individual cell segmentation compared to the Base SAM2, the fine-tuned SAM2, and the optimized SAM2.

Indeed, performance clearly improved as shown in the representative output images (Figure 3a, last column) and further confirmed in the scores (Figure 3c): from 0.184 to 0.586 (mean Jaccard) and from 0.275 to 0.718 (mean Dice) for aggregate cell segmentation compared to the Base SAM2 with default parameters. This also corresponds to a 5.6% increase in the Dice score compared to the optimized Fine-Tuned SAM2, a 705.2% increase in the Dice score compared to the fine-tuned SAM2 with default parameters, a 11.6% increase in the Dice score compared to the Base SAM2 with optimized parameters, and a 161.6% increase in the Dice score compared to the Base SAM2 with default parameters. The relative increases in performance with our lightweight pipeline were mirrored with the Jaccard indices.

Beyond numerical gains, the refined SAM2 directly addressed a major challenge in instance segmentation: reducing spurious overlapping masks that inflate aggregate coverage but misrepresent individual cells. Taken together, this case study demonstrates that rigorous instance-level segmentation is attainable with our pipeline that is both lightweight and accessible, lowering the computational and data barriers that have limited the broader adoption of foundation models in microscopy. Having established the efficacy of our lightweight pipeline in improving segmentation accuracy relative to the Base SAM2, we next evaluate our lightweight pipeline against leading segmentation tools.

### Robust benchmarking of lightweight pipeline

To study whether our lightweight pipeline can outperform widely used segmentation tools, we benchmark our refined SAM2 with several open-source pipelines and commercial standards on the identical 25-image CHO test set. Because many segmentation tools are designed primarily to recover aggregate cell area, we first compare the aggregate area Dice scores and Jaccard indices with CellProfiler, Cellpose, Imaris, ilastik (pixel classification workflow), and Otsu thresholding. We further evaluate segmentation accuracy with CellSAM, a recent segmentation tool that fine-tunes the SAM. For the tools that segment individual cells in the standard workflows (CellProfiler, CellSAM, Cellpose), we also conduct instance level Dice and Jaccard evaluations by averaging across corresponding masks with the manual segmentation (see the Methods for more details). Figure 4a shows segmentation outputs of representative images by all the methods, in addition to the original.

**Figure 4.**
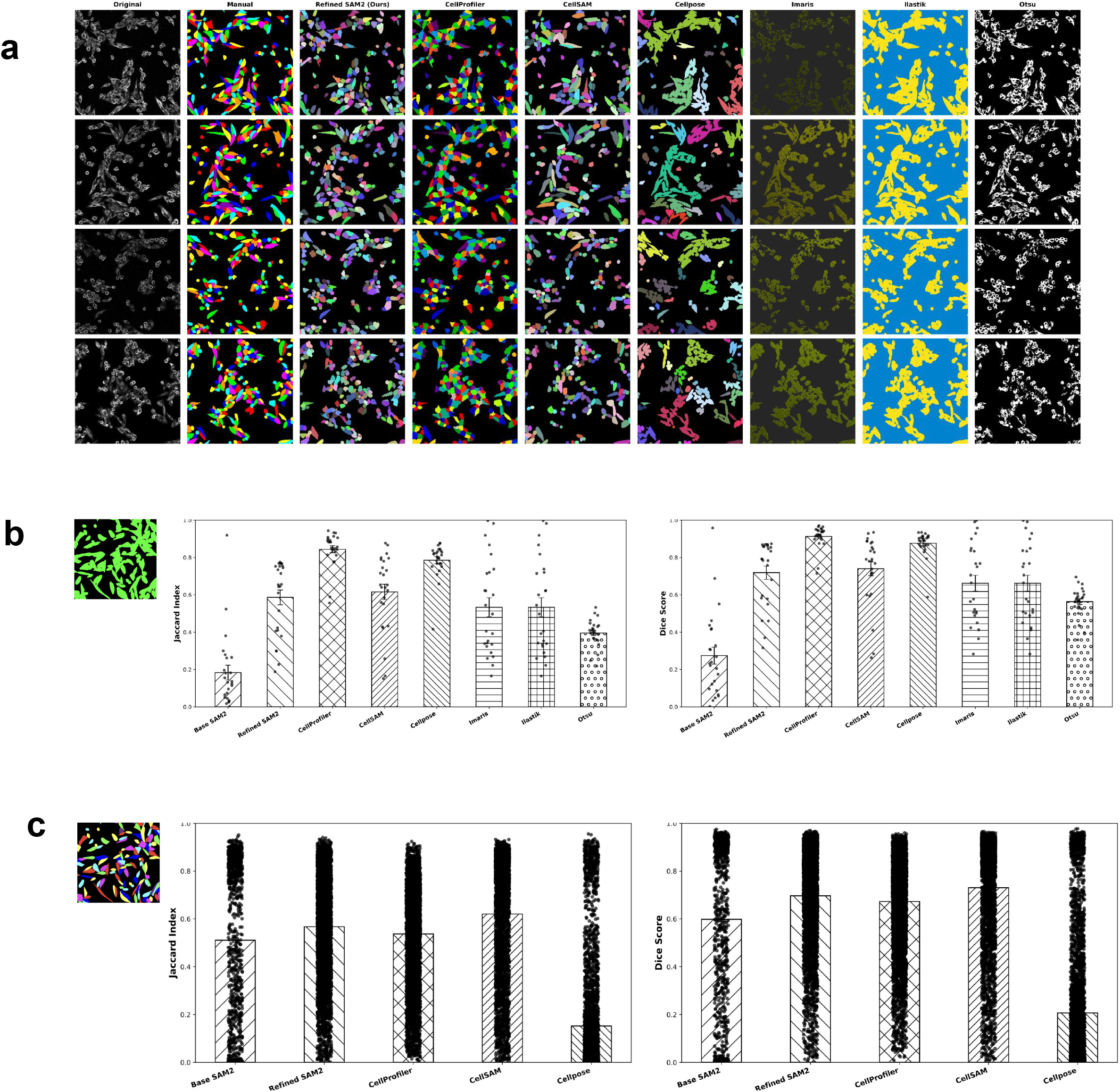
Lightweight Google Colab pipeline delivers matches or outperforms leading image segmentation software. To benchmark our lightweight pipeline against the current paradigms of cell segmentation, we compare segmentation accuracy for the same CHO cell test set used for our refined SAM2 across multiple software tools. a) Representative outputs from open-source pipeline and commercial software, compared to the original images. Dice scores and Jaccard indices for b) aggregate mask area and c) individual cell areas were measured against the manually generated ground truth for each segmentation method. Error bars represent standard error of the mean across independent image samples.

We expected segmentation accuracy to be comparable, if not exceed, existing software. Indeed, the aggregate area Dice scores shown in Figure 4b demonstrate that our refined SAM2 (0.7184), outperforms Imaris (0.6620), ilastik (0.6620), and Otsu thresholding (0.5634). Although our refined SAM2 Dice score is 2.85% less than that of CellSAM, the difference is non-significant, and CellSAM exhibits higher variability. The Dice score of refined SAM2 is 21.26% less than that of CellProfiler and 18.01% less than that of CellPose. The general trends are mirrored by the Jaccard indices. These results indicate that our lightweight pipeline can achieve high-quality aggregate segmentation results, similar to widely used segmentation tools, with substantially reduced training data and computational cost.

For the instance level evaluations, we expected our segmentation accuracy to also be comparable to existing software, given that our Hoechst-based post-processing strictly filters masks with one-to-one mappings between nuclei and cells. As predicted, our refined SAM2 demonstrated a 236.47% increase in Dice Score compared to Cellpose (p< 0.001), while performing comparably, though slightly lower, than CellProfiler (p>0.001) and CellSAM (p>0.001) (Figure 4c).

Our results demonstrate that modest fine-tuning of SAM2, paired with biologically informed post-processing, enables high-accuracy segmentation across both tissue- and cell-level microscopy tasks. The lightweight pipeline consistently improved Dice and Jaccard scores relative to Base SAM2 and achieved comparable or superior performance to leading segmentation tools, despite requiring minimal data and computation. By leveraging Google Colab’s accessible infrastructure, our framework eliminates major barriers to adapting SAM2 for microscopy. Together, these findings establish a scalable and open-source strategy for democratizing foundation model fine-tuning in biological imaging.

## Discussion

Current methods of adapting foundation models to domain-specific applications are largely inaccessible because of their reliance upon large datasets (impractical for individual laboratories to collect) and significant computational cost. To address this problem, we have developed a lightweight Google Colab pipeline for open-source SAM2 fine-tuning with compact datasets. Our pipeline involves a novel strategy of partial model fine-tuning, paired with biologically informed post-processing of model outputs. We demonstrate high accuracy across image domains without specialized hardware or modifications to the SAM2 architecture, establishing the paradigm for accessible fine-tuning of foundation models. Our cell segmentation case study establishes a fully automated method to fine-tune the SAM2 outputs for both whole-field and single-cell segmentation tasks, utilizing CellProfiler to generate training data. Our cell segmentation case study demonstrates benchmark rigor for instance segmentation, outperforming widely used cell segmentation tools. We anticipate future work to validate our lightweight pipeline across a broader range of imaging modalities and biological samples, and generate robust, fully automatic fine-tuning pipelines for H&E, brightfield, multiphoton, and electron microscopy, as well as MRI of various tissue and cell types.

We validate the improved performance of our lightweight pipeline relative to the Base SAM2 and preliminarily fine-tuned SAM2 iterations, observing consistent gains in Dice and Jaccard indices (relative to ground truth) across all segmentation tasks on held-out test sets. Although the brain segmentation case study demonstrated four images that were outliers and scored less than a 0.5 on the Jaccard index, the low scores are primarily because of the poor image quality, including weak signal and large noise.

Our cell segmentation case study demonstrates clear advantages over ilastik, Imaris, and Otsu thresholding, demonstrating higher aggregate-mask segmentation accuracy. In addition, our refined model outperforms both CellProfiler and Cellpose at the task of individual cell segmentation. Eminently, our lightweight pipeline produces instance-level masks for single-cell segmentation without requiring manual boundary annotations, thereby reducing the most labor-intensive barrier for many laboratories. This provides another clear advantage over tools such as ilastik’s Pixel Classification or Imaris’s standard workflows, which either focus on pixel-level classification or rely heavily on user input for boundary refinement. For methods that natively generate instance-level masks, our approach still offers key benefits: our model outperforms

Cellpose on the task of individual cell segmentation, and, unlike CellSAM, it remains fully open and re-trainable with a smaller computational footprint. Although we attempted to benchmark our refined model against μSAM, we were unable to adapt the published implementation to our 1024 × 1024 microscopy images, suggesting hidden dependencies on small image sizes.

Importantly, we do not assert universal superiority over these existing tools, but rather demonstrate that our pipeline offers greater accessibility and scalability while providing unique advantages, including outperforming several widely used standards and achieving comparable performance to others. The accessibility of our Google Colab pipeline establishes a practical framework for researchers to adapt the SAM2 to their own imaging domains at low computational cost.

While our contribution establishes an accessible pipeline for the domain-specific fine-tuning of SAM2, it has several limitations compared to existing methods of segmentation. Most importantly, the modest size of the training data functions as both a strength and a weakness: although it makes SAM2 fine-tuning accessible to researchers, the fine-tuning may not generalize to all edge cases. In addition, the reliance of our pipeline on biologically informed post-processing involves domain specific algorithms that do not generalize universally. However, this pipeline can be easily tailored to unique features of individual biological samples using similar algorithms for post-processing, as long as they are applied equally to the whole dataset without observer bias. Furthermore, the benchmarking was conducted with the standard workflows of each software, as opposed to exhaustive optimization. Finally, although the SAM2 architecture enables precise segmentation of objects in videos, our lightweight pipeline was not tested on microscopy video footage, which remains an important direction for future work.

Next steps to further improve our lightweight pipeline include determination of the optimal training size and testing for microscopy time-lapse videos. In the near future, we plan to automate pipelines for a broader range of microscopy domains and tasks: for example, using brain-scale fine-tuning to segment large anatomical regions, while pairing cell-level fine-tuning with biologically informed algorithms to distinguish between specific cell types such as neurons and glia, and even subtypes of neurons and glia using cell-type/stage markers. Overall, our study demonstrates the ability of modest fine-tuning of SAM2, combined with biologically informed post-processing, to deliver high-quality segmentation across diverse microscopy tasks. By lowering computational barriers, our lightweight pipeline makes foundation model adaptation accessible to more laboratories, accelerating discoveries across microscopy-based research for both fundamental biology and clinical pathology/radiology.

## Methods

### Mouse brain preparation and staining

The mice were perfused transaortically with PBS, and the brain was fixed with 4% paraformaldehyde for 24 hours, then transferred to 15% sucrose, 30% sucrose and 30% sucrose with 0.1% azide consecutively. The preserved portion was sectioned coronally with a microtome at 40um. The brain sections were immunostained with Alexa Fluor 647 anti-Tubulin β3 (0.5μg/ml, BioLegend, Cat# 801210) at 4C, overnight, and Hoechst (1μg/ml, Thermofisher, Hoechst 33342) at room temperature, 5min. Brain sections were imaged using Hamamatsu Nanozoomer s60 at 20X.

### Cell culture and staining (CHO-K1 cells)

CHO-K1 Cells (CCL-61) were grown in DMEM/F12 (Life Technologies 11320082) supplemented with FBS and pen/strep (Life Technologies 15070-063). Cells were plated on 8 well chamber slides (Life Technologies 154941) and then fixed with 4% paraformaldehyde.

Cells were permeabilized with 0.25% Triton-X in TBS for 15 minutes then blocked with 10% NGS in 0.02% TritonX-100 in TBS for 1 hr at room temperature. Cells were stained with Alexa Fluor 488 Anti-alpha Tubulin antibody (0.5μg/ml, Abcam, ab195887) at 4C, overnight, and Hoechst (1μg/ml, Thermofisher, Hoechst 33342) at room temperature, 5min. Cells were imaged using Olympus laser scanning confocal at 40X.

### Image acquisition

Brain images were acquired using the Hamamatsu Nanozoomer s60 with an NDPIS file format. The slides were imaged at 20X magnification with a resolution of 441 nm/pixel using the DAPI, FITC, TRITC, and Cy5 fluorescence filters. The exposure time was adjusted and optimized for each sample ranging from 80ms to 250ms to account for over saturation. Only 405 (Hoechst) and 647 (Tubulin) images were used for this study.

The cell images were acquired using the Olympus FV31S-SW with an OIR file format. The slides were imaged at 40X using the UPLSAPO 40X objective and a numerical aperture (NA) of 0.95. The images were acquired at a resolution of 512*512 pixels, using Galvano scan mode in one-way direction, line sequential, at a sampling speed of 8 us/pixel. The channels used were DM405 nm for Hoechst, 488 nm for Alexa Fluor 488, 561 nm for Alexa Fluor 568, and 640 nm for Alexa Fluor 647. The laser intensity (%), sensitivity (HV), and the gain and offset were optimized for the images to balance signal to noise ratio and prevent over saturation. Only 405 (Hoechst) and 488 (Tubulin) images were used for this study.

### Dataset preparation

Hoechst-stained mouse brain images were annotated by hand to generate ground truth segmentations of the hippocampus. Cell image ground truth segmentations were generated with the CellProfiler pipeline, which identified and outlined individual CHO cells from the tubulin channel. See *CellProfiler* below for more details. Raw cell images in OIR format were converted to 8-bit TIFF files, and the tubulin and Hoechst channels were extracted for downstream processing. Each channel underwent automatic contrast adjustment^8^ in FIJI (ImageJ v3.0) to normalize intensity distributions across images.

### SAM2 fine-Tuning

We fine-tuned the SAM-2.1^19^ hiera-base plus model, which consists of a hierarchical vision transformer image encoder, a prompt encoder for points, boxes, and text, and a lightweight transformer-based mask decoder. The mask decoder was trained end-to-end, while the prompt and image encoders remained frozen to preserve general prompting ability. Training images were resized to 1024 × 1024 resolution, and optimization was performed with AdamW^26^ using cosine annealing^27^ over 40 epochs. A batch size of 1 was used due to memory constraints, with learning rates set to 5.0 × 10^−6^ for the base model and 3.0 × 10^-6 for vision-specific components. The loss function combined Dice loss^28^ and binary cross-entropy^29^ in a 2:1 ratio. Training was conducted on a NVIDIA A100 GPU (80 GB).

### Parameter optimization

SAM2 provides a range of advanced tunable parameters that govern mask generation beyond the learned weights of the model^19^. Together, these parameters determine how densely the model explores potential masks and how aggressively it filters them. To maximize segmentation performance, joint hyperparameter optimization was done on the validation set. Rather than tuning parameters independently, multiple hyperparameters were varied simultaneously to account for potential interaction effects. The search space included sampling density parameters (points_per_side, points_per_batch), post-processing thresholds (pred_iou_thresh, box_nms_thresh), stability-based filtering parameters (stability_score_thresh, stability_score_offset), hierarchical refinement depth (crop_n_layers), and constraints on the minimum region size (min_mask_region_area). Model performance was evaluated on the validation set using the Jaccard Index and Dice coefficient as primary metrics.

### Brain post-optimization algorithm

For brain tissue images, the hippocampal region was consistently present among the three highest-scoring masks according to the model’s internal confidence estimates. To reliably identify the correct region, we applied an intensity-based heuristic: the mask with the highest proportion of white pixels was selected, reflecting the strong Hoechst fluorescence associated with the high density of dentate gyrus in the hippocampus.

### Cell post-optimization algorithm

For single-cell images, additional refinement was required to ensure accurate and non-redundant cell boundaries. Raw Hoechst channel often displayed a diffuse gray background that obscured nuclear structures. To reduce this interference, we suppressed low-intensity pixels by applying an intensity threshold of 50 (on a scale of 0–255) to the 405 channel. In addition, masks in which more than 50% of pixels were black were then discarded, ensuring that only regions with sufficient nuclear signal were kept for downstream analysis.

To minimize redundancy, overlapping masks were compared in pairs. When two masks shared more than 80% of the smaller mask’s area, only the higher-quality mask was retained. This step reduced duplication and ensured that each nucleus was represented by a single boundary.

The center of each nucleus was then recorded from the remaining masks. These points served as anchors to enforce a one-to-one relationship between nuclei and cell masks. Candidate masks were considered valid if they contained the center of a nucleus, and only one mask was assigned per nucleus. In cases where multiple candidates were possible, the mask with the largest area was chosen to maximize boundary coverage.

### Evaluation metrics (Dice, Jaccard)

Segmentation performance was assessed using the Dice Coefficient^23^ and the Jaccard Index^24^. For the brain images these were computed relative to the manually generated ground truths. The Dice Coefficient was calculated as 2|A ∩ B| /(|A| + |B|). This measure is widely used in medical image segmentation and is particularly suited for evaluating smaller structures. The Jaccard Index was calculated as |A ∩ B| /|A ∪B|. This metric provides a measure of spatial agreement and is especially sensitive to how well boundaries are captured.

For single-cell segmentation tasks, Dice and Jaccard scores were computed for each individual cell mask and then averaged across all cells within an image. For aggregate segmentation tasks, scores were computed using the union of all predicted masks compared with the union of all ground-truth regions.

### Cellpose

We benchmarked our lightweight pipeline against Cellpose^15^ (v4.0.1+), running inference with the Cellpose “cyto” model with automatic diameter estimation (v4 API). Model outputs were post-processed to remove fragments smaller than 50 pixels and converted to labeled instance masks.

### CellSAM

CellSAM^20^ (v1.2) was run with default parameters and pretrained weights. Inference was performed through the segment_cellular_image API on a single GPU (CUDA backend). Each test image was processed to generate an instance label map and a stack of binary masks, corresponding to individual segmented objects. Following segmentation, duplicate labels and background regions were excluded.

### Imaris

Cell masks for the confocal images were generated through Imaris^9^ using the Surface Creator tool to define cell boundaries based on set uniform intensity thresholds. All surfaces were reconstructed to capture the majority of the cell morphology while minimizing background signal. Object–object statistics were applied to quantify spatial relationships between segmented cells and ensured that the segmentation was done by contrasting the background with the cells themselves. This approach provided consistent segmentation across the dataset while preserving biologically and morphologically cell features that were both significant and clear from the perspective of the Imaris segmentation program.

### Ilastik

Cell segmentation of the original confocal images was performed using the machine learning–based pixel classification framework implemented in Ilastik^16^. To optimize boundary delineation and minimize background interference, all available feature categories—including color, edge, and texture descriptors—were incorporated into the classification process. For system training, representative cells were manually annotated in yellow, while background regions were annotated in blue, thereby enabling the classifier to distinguish cellular structures from non-cellular signals. Following training, Ilastik extracted and generalized these features to the complete image set, applying consistent classification criteria across all samples. By enabling both the “Live Update” and “Segmentation” functions, the pixel classification workflow produced reproducible cell segmentations against their respective backgrounds.

### CellProfiler

The images were split by the channels, uploaded to CellProfiler^17^, then organized by metadata, name and type. The Hoechst channel was used to identify the cell nuclei as the primary object and this segmented the nuclei. The tubulin channel was used to identify the mask being used and set as the secondary object to detect the cell boundaries. Following these identifications and assignments, the outlines of the secondary objects were overlayed on the tubulin image and saved in grayscale. The objects were then converted to images, masked using the secondary objects identified (cell), inverted to remove any background, and saved as an image for exporting. The saved outline and masks were processed further and applied in the subsequent phases.

### Statistical analysis and reproducibility

For hippocampus segmentation, 54 Hoechst-stained mouse brain images with manually annotated hippocampal masks were used for training, and 24 images were held out for testing. For single-cell segmentation, 59 tubulin-stained CHO cell images with CellProfiler-generated masks were used for training, and 25 images were held out for testing.

## Data Availability

https://drive.google.com/drive/folders/1YO3DPqXuL8WpTpjOxuhtRFa8oWqWohrN?usp=sharing

### Code Availability

https://github.com/sharvaa-selvan/SAM2-Fine-Tuning

## Author Contributions

E.B., S.S., M.S., Y.S. conceptualized the work. S.O., Z.D., V.L., M.N. provided experimental methods, resources and data. E.B., S.S., Y.S. wrote and edited the manuscript. All authors reviewed and approved the final manuscript.

## Acknowledgements

We thank Massachusetts General Institute for Neurodegenerative Disease (MIND), the Department of Neurology at MGB, and the Neuroscience Institute (NSI) for support of the Song Laboratory. We acknowledge Meta AI for developing the Segment Anything Model 2 (SAM2), which served as the foundation for our fine-tuning and post-processing pipeline. Computational analyses were conducted using Google Colab open-source resources.

## Funding

The authors would like to acknowledge all the funding resources: NIH AG072516, Jack Satter Award, Pape Adams ALS Transformative Scholar Award, AARG grant from the Alzheimer’s Association, Healey Center ALS Young Scholar Award, and Mass General Hospital Mussallem Transformative Scholar Award in ALS Research to YS.

## Notes

### Competing Interest Statement

The authors have declared no competing interest.

## References

1 Meijering, E. Cell segmentation: 50 years down the road. IEEE Signal Process Mag. 29, 140–145 (2012).

2 Paddock, S. W. e. Confocal Microscopy: Methods and Protocols. (Springer, 2014).

3 Kumar, R. K. et al. Virtual microscopy for learning and assessment in pathology. J Pathol 204, 613–618, doi:10.1002/path.1658 (2004).

4 Aeffner, F. et al. Digital Microscopy, Image Analysis, and Virtual Slide Repository. ILAR J 59, 66–79, doi:10.1093/ilar/ily007 (2018).

5 Zuraw, A. & Aeffner, F. Whole-slide imaging, tissue image analysis, and artificial intelligence in veterinary pathology: An updated introduction and review. Vet Pathol 59, 6–25, doi:10.1177/03009858211040484 (2022).

6 Yu, Y. W. C.; Fu, Q.; Kou, R.; Huang, F.; Yang, B.; Yang, T.; Gao, M. Techniques and challenges of image segmentation: a review. Electronics 12 (2023).

7 Schindelin, J. et al. Fiji: an open-source platform for biological-image analysis. Nat Methods 9, 676–682, doi:10.1038/nmeth.2019 (2012).

8 Broeke, J., Mateos Pérez, J. M. & Pascau, J. 58 (Packt Publishing, 2015).

9 Imaris 10.2 (Oxford Instruments, Zürich, Switzerland 2024).

10 Yushkevich, P. A. et al. User-guided 3D active contour segmentation of anatomical structures: significantly improved efficiency and reliability. Neuroimage 31, 1116–1128, doi:10.1016/j.neuroimage.2006.01.015 (2006).

11 Yushkevich, P. A., Yang, G. & Gerig, G. ITK-SNAP: An interactive tool for semiautomatic segmentation of multi-modality biomedical images. Annu Int Conf IEEE Eng Med Biol Soc 2016, 3342–3345, doi:10.1109/EMBC.2016.7591443 (2016).

12 Dimopoulos, S., Mayer, C. E., Rudolf, F. & Stelling, J. Accurate cell segmentation in microscopy images using membrane patterns. Bioinformatics 30, 2644–2651, doi:10.1093/bioinformatics/btu302 (2014).

13 Peng, H. Bioimage informatics: a new area of engineering biology. Bioinformatics 24, 1827–1836, doi:10.1093/bioinformatics/btn346 (2008).

14 Stringer, C. & Pachitariu, M. Cellpose3: one-click image restoration for improved cellular segmentation. Nat Methods 22, 592–599, doi:10.1038/s41592-025-02595-5 (2025).

15 Stringer, C., Wang, T., Michaelos, M. & Pachitariu, M. Cellpose: a generalist algorithm for cellular segmentation. Nat Methods 18, 100–106, doi:10.1038/s41592-020-01018-x (2021).

16 Berg, S. et al. ilastik: interactive machine learning for (bio)image analysis. Nat Methods 16, 1226–1232, doi:10.1038/s41592-019-0582-9 (2019).

17 Carpenter, A. E. et al. CellProfiler: image analysis software for identifying and quantifying cell phenotypes. Genome Biol 7, R100, doi:10.1186/gb-2006-7-10-r100 (2006).

18 Kirillov, A. M., E.; Ravi, N.; Mao, H.; Rolland, C.; Gustafson, L.; Xiao, T.; Whitehead, S.; Berg, A. C.; Lo, W. Y.; Dollár, P.; Girshick, R. Segment Anything. arXiv:2304.02643 (2023).

19 Ravi, N. G. V.; Hu, Y.-T.; Hu, R.; Ryali, C.; Ma, T.; Khedr, H.; Rädle, R.; Rolland, C.; Gustafson, L.; Mintun, E.; Pan, J.; Alwala, K. V.; Carion, N.; Wu, C.-Y.; Girshick, R.; Dollár, P.; Feichtenhofer, C. Segment Anything Model 2 (SAM 2): segment anything in images and videos. arXiv:2408.00714 (2024).

20 Israel, U. et al. CellSAM: A Foundation Model for Cell Segmentation. bioRxiv, doi:10.1101/2023.11.17.567630 (2025).

21 Archit, A. et al. Author Correction: Segment Anything for Microscopy. Nat Methods 22, 1603, doi:10.1038/s41592-025-02745-9 (2025).

22 Jackson, D. A. S. K. M.; Harvey, H. H. Similarity Coefficients: Measures of Co-Occurrence and Association or Simply Measures of Occurrence? The American Naturalist 133, 436–453 (1989).

23 Dice, L. R. Measures of the amount of ecologic association between species. Ecology 26, 297–302 (1945).

24 Jaccard, P. Étude comparative de la distribution florale dans une portion des Alpes et des Jura. Bulletin de la Societe Vaudoise des Sciences Naturelles 37, 547–579 (1901).

25 Caicedo, J. C. et al. Publisher Correction: Nucleus segmentation across imaging experiments: the 2018 Data Science Bowl. Nat Methods 17, 241, doi:10.1038/s41592-020-0733-z (2020).

26 Loshchilov, I. H. F. Decoupled Weight Decay Regularization. arXiv:1711.05101 (2017).

27 Loshchilov, I. H. F. SGDR: Stochastic gradient descent with warm restarts. arXiv:1608.03983 (2016).

28 Sudre, C. H., Li, W., Vercauteren, T., Ourselin, S. & Jorge Cardoso, M. Generalised Dice Overlap as a Deep Learning Loss Function for Highly Unbalanced Segmentations. Deep Learn Med Image Anal Multimodal Learn Clin Decis Support (2017) 2017, 240–248, doi:10.1007/978-3-319-67558-9_28 (2017).

29 Ruby, U. Y. V. Binary cross entropy with deep learning technique for image classification. International Journal of Advanced Trends in Computer Science and Engineering 9.10 (2020).

